# Glyoxal fixation facilitates transcriptome analysis after antigen staining and cell sorting by flow cytometry

**DOI:** 10.1101/2020.10.05.326082

**Authors:** Prasanna Channathodiyil, Jonathan Houseley

## Abstract

A simple method for extraction of high quality RNA from cells that have been fixed, stained and sorted by flow cytometry would allow routine transcriptome analysis of highly purified cell populations and single cells. However, formaldehyde fixation impairs RNA extraction and inhibits RNA amplification. Here we show that good quality RNA can be readily extracted from stained and sorted mammalian cells if formaldehyde is replaced by glyoxal - a well-characterised fixative that is widely compatible with immunofluorescent staining methods. Although both formaldehyde and glyoxal efficiently form protein-protein crosslinks, glyoxal does not crosslink RNA to proteins nor form stable RNA adducts, ensuring that RNA remains accessible and amenable to enzymatic manipulation after glyoxal fixation. We find that RNA integrity is maintained through glyoxal fixation, permeabilisation with methanol or saponin, indirect immunofluorescent staining and flow sorting. RNA can then be extracted by standard methods and processed into RNA-seq libraries using commercial kits; mRNA abundances measured by poly(A)+ RNA-seq correlate well between freshly harvested cells and fixed, stained and sorted cells. We validate the applicability of this approach to flow cytometry by staining MCF-7 cells for the intracellular G2/M-specific antigen cyclin B1 (CCNB1), and show strong enrichment for G2/M-phase cells based on transcriptomic data. Switching to glyoxal fixation with RNA-compatible staining methods requires only minor adjustments of most existing staining and sorting protocols, and should facilitate routine transcriptomic analysis of sorted cells.

## Introduction

High-throughput RNA sequencing (RNA-seq) is increasingly used as a first-pass method to characterise cell populations, providing detailed and robust data through standardised pipelines. In parallel, purification of cell populations by flow cytometric cell sorting has become routine, with the capacity to isolate cells based on simultaneous quantification of many antigens. It would seem natural to unite these techniques, such that carefully purified cell populations are characterised by RNA-seq as standard, however this has proved surprisingly problematic primarily due to difficulties in RNA recovery from fixed and stained cells.

Ideally, cells destined for RNA-seq analysis should be fixed before removal from growth media or tissue context as tissue dissociation and cell staining methods induce stress responses that affect the transcriptome [1-3]. Formaldehyde is the fixative of choice for the vast majority staining protocols but, as pathologists are well aware, recovery of intact RNA from formaldehyde-fixed material is not trivial [4-6]. Various labs have successfully performed transcriptome analysis of formaldehyde fixed and sorted cells using methods such as MARIS and FIN-seq [7-10], but these require specialised RNA extraction and de-crosslinking procedures that are not compatible with many downstream applications.

Although RNA recovered from formaldehyde-fixed cells is often of low quality, formaldehyde is a standard reagent in RNA electrophoresis and does not cause RNA degradation of itself [11]. Instead, problems arise firstly from RNA-protein crosslinking: formaldehyde reacts with the primary amine of guanine, adenine or cytosine to form a Schiff base that can be attacked by nucleophiles such as lysine residues of nearby proteins to create stable covalent crosslinks [12] (Fig. 1A). This outcome is ideal when stable nucleic acid-protein complexes are required, for example in chromatin immunoprecipitation, but is very problematic when RNA needs to be purified away from proteins. If RNA extraction from formaldehyde-fixed cells is attempted with widely used guanidine thiocyanate (GTC)-phenol reagents (TRI Reagent, TRIzol, etc.) then the crosslinked RNA-protein complexes partition to the interphase and are lost on phase separation [12, 13]. Proteinase K treatment can be used to deproteinise and extract RNA from formaldehyde-fixed material [14], however covalent crosslinks must still be reversed by heat treatment to remove adducts that otherwise inhibit enzymatic reactions: peptides remain attached after proteinase K digestion, and even once these are removed polymerases can still be inhibited by residual methylol groups, accounting for the poor amplification of DNA and RNA from formaldehyde-fixed material [13, 15, 16]. Furthermore, these residual adducts inhibit base-pairing and so will certainly impede purification of poly(A)+ RNA by binding to oligo(dT), a key step in many bulk and single-cell mRNA-seq protocols [17, 18]. Although crosslinks and residual adducts can be removed by heating, balancing sufficient crosslink reversal against thermal degradation of RNA is challenging [13, 19, 20].

**Figure 1:**
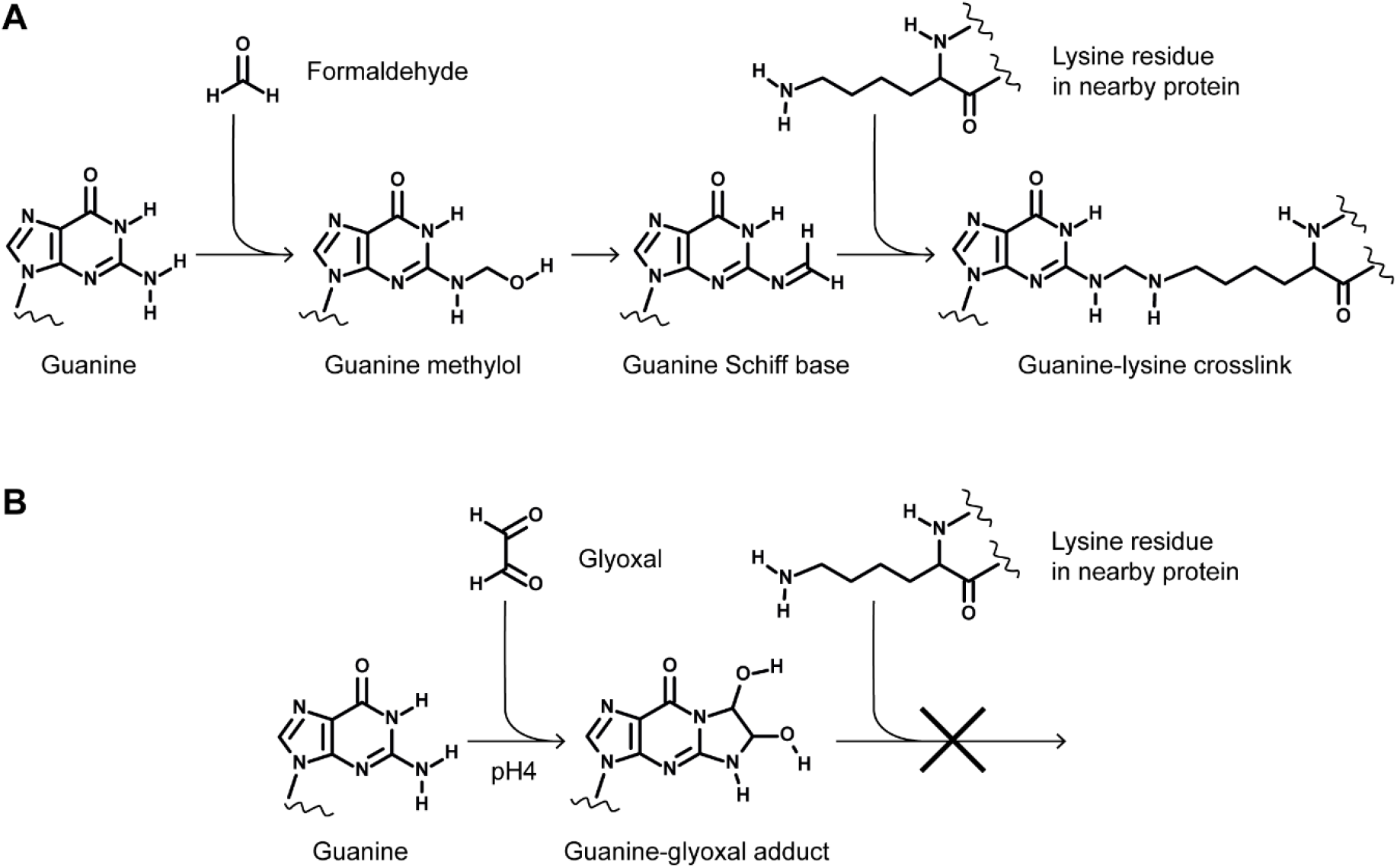
Schematic representation of formaldehyde and glyoxal reactions with guanine nucleotides. **A**: Reaction of formaldehyde with guanine, equivalent reactions occur for adenine and cytidine. **B**: Reaction of glyoxal with guanine, reaction products with other bases are not stable [31].

Of course, other fixatives are known that do not crosslink RNA to protein or introduce stable adducts. We and others routinely employ ethanol fixation on yeast prior to RNA isolation and find the fixation speed and preservation of RNA to be excellent [21, 22], while similarly high quality RNA has been isolated from methanol or ethanol-fixed mammalian cells [23-26]. Another option is glyoxal, a protein crosslinking fixative that performs similarly to formaldehyde in most applications [27-29]. Glyoxal, like formaldehyde, has long been used as an RNA denaturant in gel electrophoresis so it is known to maintain RNA integrity [30], and has useful characteristics including reduced toxicity and increased stability in solution. Importantly, although glyoxal forms protein-protein crosslinks with a similar efficiency to formaldehyde, glyoxal reacts very differently with nucleic acids; only guanine reacts to any measurable extent with glyoxal and in doing so creates a stable heterocycle that does not form covalent crosslinks with proteins under normal conditions [31] (Fig. 1B). Furthermore, the guanine-glyoxal heterocycle is unstable at pH>7 and so inhibitory adducts rapidly dissociate under standard buffer conditions [31].

Non-RNA-protein crosslinking fixatives therefore have the potential to be extremely useful in transcriptomic studies of flow sorted cells. However, little is known about the survival of RNA in alcohol fixed cells during downstream staining and sorting procedures, and to our knowledge the survival of RNA in glyoxal fixed cells has not been investigated. Here we show that glyoxal fixation allows extraction of high-quality RNA using standard protocols from stained and sorted cell samples, and that this RNA can be processed into high quality RNA-seq libraries.

## Results

### Optimisation of staining steps for RNA extraction

To determine the effectiveness of ethanol fixation in preserving intact and accessible RNA in mammalian cells, we extracted RNA from COLO205 cells that were either unfixed, fixed with 70% ethanol or fixed with 4% formaldehyde. RNA extraction was performed using TRI reagent, a monophasic GTC-phenol RNA extraction solution, and 20% of the RNA obtained was analysed on denaturing RNA mini-gels. Plentiful high quality RNA was obtained from the unfixed cells, whereas we did not recover detectable RNA from formaldehyde-fixed cells which was as expected given that proteinase K digestion was not performed (Fig. 2A, compare lanes 1 and 3). A high yield of total RNA was also obtained from ethanol fixed cells but this RNA was partially degraded (Fig. 2A, compare lanes 1 and 2), and similar problems were observed with 100% methanol fixation (Fig. 2B, lanes 1-3). We suspect that degradation occurs because alcohols permeabilise membranes and allow extracellular and intra-organellar ribonucleases to enter the cytoplasm and degrade RNA before these enzymes are denatured. Whatever the reason, this partial degradation was problematic as the extent of degradation varied between experiments, and is also likely to be variable between cell types and tissue samples depending on local and intracellular RNase concentrations.

**Figure 2:**
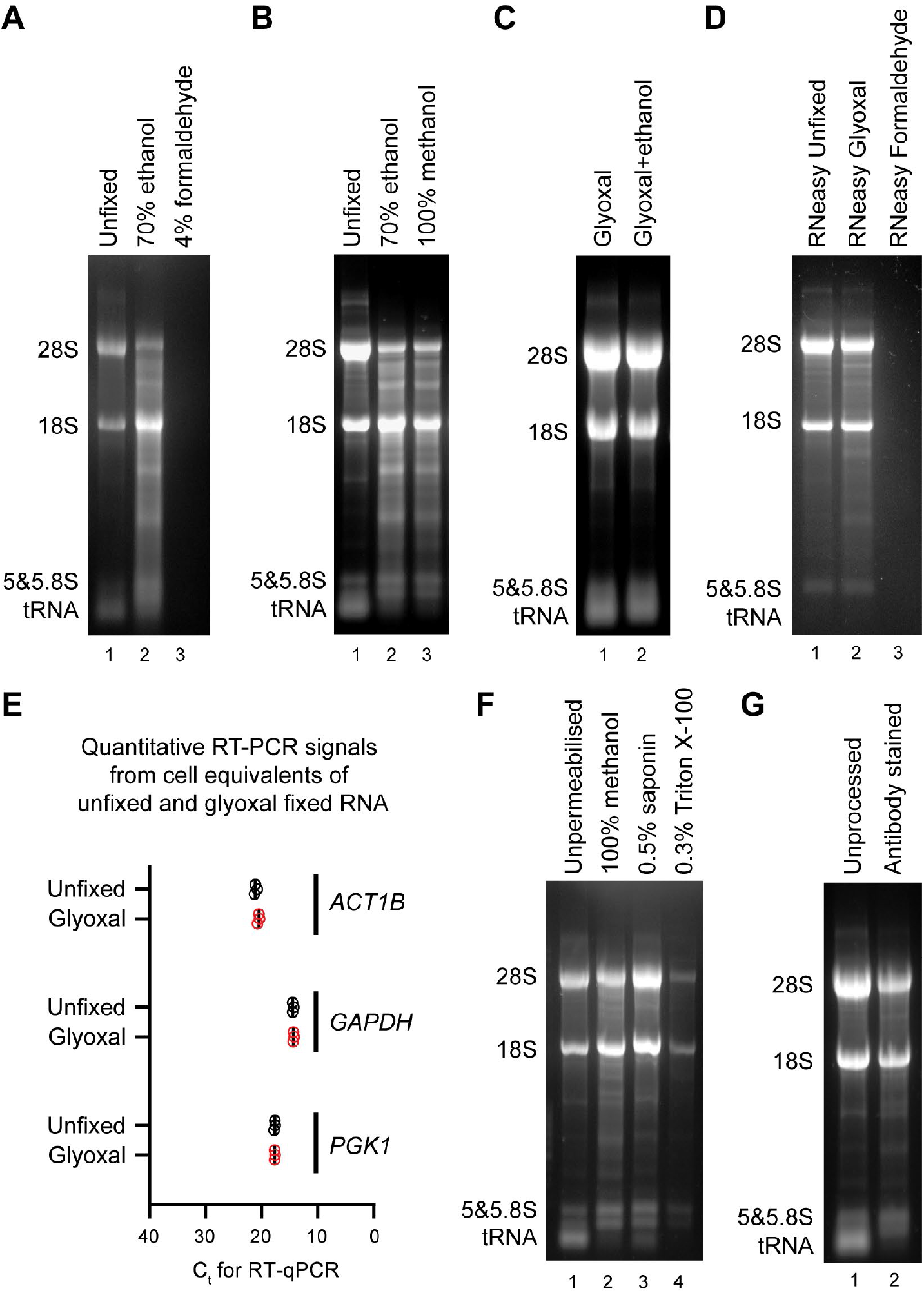
Determination of RNA-compatible fixation and permeabilisation conditions. **A**: 1×10^6^ COLO205 cells were either unfixed (lane 1), fixed with 70% ethanol on ice for 15 minutes (lane 2) or fixed with 4% formaldehyde on ice for 15 minutes (lane 3). Unfixed cells were dissolved immediately in TRI Reagent, fixed cells were washed once in PBS by centrifugation at 2000 x *g* for 3 minutes at 4°C before RNA extraction with TRI Reagent. 20% of RNA obtained was separated on a 1.2% glyoxal gel and imaged by ethidium bromide staining. **B**: COLO205 cells were either unfixed (lane 1), fixed with 70% ethanol on ice for 15 minutes (lane 2) or fixed with 100% methanol on ice for 15 minutes (lane 3). Unfixed cells were dissolved immediately in TRI Reagent, fixed cells were washed once in PBS by centrifugation at 2000 x *g* for 3 minutes at 4°C before RNA extraction with TRI Reagent. RNA was analysed as in **A**. **C**: COLO205 cells were fixed with glyoxal fixation mix (pH4) either without or with 20% ethanol and incubated on ice for 15 minutes and washed once in PBS by centrifugation at 2000 x *g* for 3 minutes at 4°C. RNA was extracted and analysed as in **A**. **D**: COLO205 cells were either unfixed (lane 1), fixed with glyoxal fixation mix (pH4) with 20% ethanol (lane 2) or with 4% formaldehyde on ice for 15 minutes (lane 3). Cells were washed once in PBS by centrifugation at 2000 x *g* for 3 minutes at 4°C and incubated on ice for 1 hour in 100 μl PBS followed by centrifugation at 2000 x *g* for 3 minutes at 4°C before RNA extraction with an RNeasy mini kit. **E**: 100 ng RNA per reaction from **D** was subjected to one-step combined reverse transcription and quantitative PCR reactions for *ACT1B, GAPDH* and *PGK1.* Ct is the cycle number at which the fluorescence exceeded threshold. 3 technical replicates for each RT-qPCR reaction were performed. **F**: COLO205 cells were either unfixed (lane 1) or fixed with glyoxal fixation mix (pH4) with 20% ethanol on ice for 15 minutes and permeabilised in 100% methanol on ice for 30 minutes (lane 2), or 0.5% saponin on ice for 30 minutes (lane 3) or 0.3% Triton X-100 on ice for 30 minutes (lane 4). RNA was analysed as in **A**. **G**: COLO205 cells were either unfixed (lane 1) or fixed with glyoxal fixation mix (pH4) with 20% ethanol on ice for 15 minutes, permeabilised in 100% methanol on ice for 30 minutes followed by incubation in primary antibody for 1 hour on ice and secondary antibody for 30 minutes on ice in dark (lane 2). RNA was analysed as in **A**.

Since alcohol fixation did not preserve RNA well in our cell-line of choice, we turned to glyoxal fixation. Two formulations of acidic 3% glyoxal fixative have recently been validated for immunostaining, either with or without 20% ethanol [27], and TRI Reagent extraction yielded RNA of excellent quality from COLO205 cells fixed with either glyoxal formulation (Fig. 2C). All further experiments were performed with glyoxal fixative containing 20% ethanol as this is reported to improve sample morphology [27]. A similar experiment involving RNA extraction with a commercial column-based kit (RNeasy, Qiagen) also yielded high quality RNA from glyoxal-fixed cells compared to unfixed cells, controlled against formaldehyde fixation which again completely impaired extraction (Fig. 2D). Although glyoxal does form adducts with guanine nucleotides, based on the measured dissociation rate of the guanine-glyoxal adduct we calculate that <1% of guanine-glyoxal adducts would remain after 3 hours at pH7.4 (the pH of PBS) [31]. This is the minimum time realistically required for immunofluorescent staining and cell sorting by flow cytometry, so RNA extracted from glyoxal-fixed, stained and sorted cells should not carry inhibitory adducts at a concentration meaningful for RNA-seq analysis. To ensure that residual adducts do not interfere with reverse transcription or PCR amplification, we performed one-step quantitative RT-PCR reactions on RNeasy-extracted RNA for large (2-300 bp) amplicons, and observed no difference between signals from unfixed and glyoxal-fixed RNA (Fig. 2E). Together these observations show that glyoxal is a superior fixative to formaldehyde or alcohols for RNA preservation in mammalian cells, and allows high quality RNA extraction by both GTC-phenol and column based methods.

One advantage of alcohol-based fixatives compared to glyoxal is the concurrent permeabilisation of cell membranes, allowing antibody ingress for staining of intracellular epitopes. We therefore tested three common permeabilisation reagents after glyoxal fixation: cold 100% methanol, 0.5% saponin and 0.3% Triton X-100 (Fig. 2F). Of these reagents, methanol and saponin both performed well with only Triton X-100 impairing RNA yield or quality; this suggests that permeabilisation by Triton X-100 but not saponin or methanol allows RNA to escape through the cell membrane. Notably, the developers of PRIMMUS, a method for intracellular staining of proteins followed by mass spectrometry, similarly concluded that Triton X-100 but not methanol permeabilisation lead to a significant loss of cellular proteins [32]. However, we noticed that the signal from small RNA species, particularly tRNA, was dramatically reduced in all permeabilised or alcohol fixed samples (Fig. 2B compare tRNA intensity in lanes 2&3 to lane 1, or Fig. 2F lanes 2&3 compared to lane 1) indicating that RNA species <100 nt in length will be lost in almost any staining procedure involving permeabilisation, and therefore that staining for intracellular antigens may not be compatible with recovery of small RNAs and miRNAs.

To confirm that high RNA quality and yield are maintained after staining, we then compared RNA from cells lysed directly in TRI Reagent after harvest against RNA from cells subjected to glyoxal fixation and methanol permeabilisation followed by a 2-step primary and secondary antibody staining procedure. The quality of RNA was equivalent between the stained and unprocessed cells and yield only slightly reduced (Fig. 2G). We therefore find that glyoxal fixation followed by methanol permeabilisation allows immunostaining of intracellular antigens and recovery of high quality RNA by standard methods without requirement for proteinase digestion or de-crosslinking.

### Glyoxal fixed cells yield high quality RNA-seq libraries after flow sorting

We chose Cyclin B1 (CCNB1) as a trial intracellular antigen for validation of RNA-seq analysis in glyoxal fixed and stained cells. CCNB1 accumulates during G2 phase of the cell cycle and is degraded at the end of M-phase, so the quality of staining and sorting based on CCNB1 can be verified by comparison to DAPI staining for DNA content. Furthermore, gene expression changes across the cell cycle are well characterised and so the successful purification of G2/M-phase cells should be detectable based on differential expression of cell cycle genes in RNA-seq data.

To determine whether our staining protocol reproducibly maintained RNA quality, we compared cells immediately dissolved in TRI Reagent at harvest to cells that underwent staining. Antibody staining was performed according to the consensus protocol given in the Materials and Methods section entailing glyoxal fixation, methanol permeabilisation and two-step immunostaining with rabbit anti-CCNB1 followed by Alexa Fluor 488-labelled donkey anti-rabbit. Two replicate experiments were performed for COLO205 cells and two for MCF-7 cells, RNA was extracted with TRI reagent and RNA quality assessed by Bioanalyzer. Representative Bioanalyzer plots are shown (Fig. 3A) and details of RNA integrity and yield are given in Table 1. The RNA integrity number (RIN), which varies from 1-10 was passable for RNA from stained COLO205 cells (>7.6) and very high for RNA from stained MCF7 cells (>9.5). Importantly, there was little reduction in RNA quality during staining (the worst outcome was a reduction in RIN of 0.9 in one COLO205 replicate). Furthermore, extractions were efficient with >40% of the RNA yield obtained from stained cells compared to directly harvested cells.

**Figure 3:**
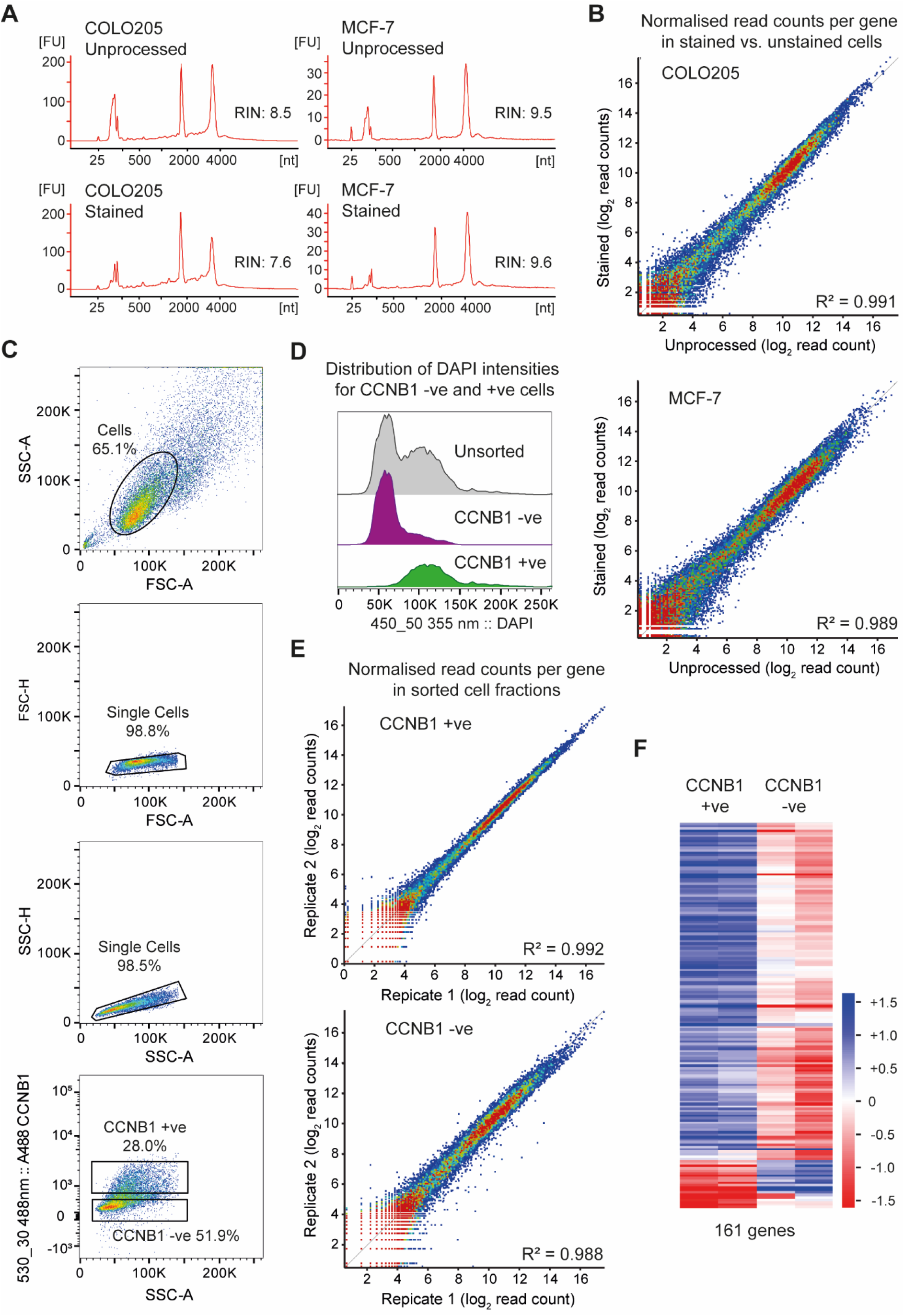
RNA-seq of stained and sorted cell populations. **A**: Bioanalyzer profiles of total RNA derived from unprocessed cells (harvested directly from trypsinised cells) compared to cells that have undergone glyoxal fixation, permeabilisation with 100% methanol and indirect immunofluorescence for CCNB1. 0.1-5 ng of total RNA was separated on total RNA pico chips on Agilent 2100 Bioanalyzer. **B**: Scatter plots comparing the normalised read count for each annotated gene in GRCh38 in poly(A)+ RNA-seq libraries derived from unprocessed and from glyoxal fixed, permeabilised and stained cells. Two biological replicates were sequenced for each condition and averaged, read counts were normalised to the total number of reads in each library. Data is shown for both COLO205 and MCF-7 cells. **C**: Flow cytometry density plots for MCF-7 cells labelled with anti-CCNB1 primary antibody and donkey Alexa Fluor-488 conjugated secondary antibody and sorted using a BD FACSAria III sorter. Intact cells (shown within the elliptical gate) were distinguished from off-scale events and cell debris using forward scatter (FSC) and side scatter (SSC) measurements (panel 1). Doublets were excluded from the gated cells by a 2-step gating strategy with pulse height (H) plotted against area (A) for FSC parameter followed by SSC-H versus SSC-A plot (panel 3). Fluorescence thresholds for isolation of CCNB1 positive and negative cell fractions are shown in panel 4. Gates were set using the negative control and the CCNB1 positive and negative sorting gates were moved apart from each other to maximise purity when sorted. **D**: Fluorescence histogram of DAPI intensities for sorted CCNB1 negative and positive populations **E**: Scatter plots comparing the normalised read count for each annotated gene in GRCh38 in poly(A)+ RNA-seq libraries derived from two biological replicates each of CCNB1 positive (left) and negative (right) MCF-7 cells. Note the particularly tight correlation after sorting towards the antigen of interest. **E**: Hierarchical clustering analysis of the 161 significantly different expressed genes identified DEseq2 analysis (p<0.05) comparing CCNB1-positive and CCNB1-negative MCF-7 samples.

**Table 1:** Details of RNA extraction and library construction.

|  | *Number of cells* | *RNA yield (ng)* | *RIN* | *Library Input*<br>*RNA (ng)* | *Concentration*<br>*of library (nM)* |
| --- | --- | --- | --- | --- | --- |
| **COLO205** |  |  |  |  |  |
| Unprocessed | 1.25x10<sup>6</sup> | 14298 | 8.5 | 20 | 15.43 |
| Stained | 1.25x10<sup>6</sup> | 13269 | 7.6 | 20 | 9.53 |
| Unprocessed | 1.25x10<sup>6</sup> | 10434 | 8.1 | 20 | 4.9 |
| Stained | 1.25x10<sup>6</sup> | 10531.5 | 7.7 | 20 | 17.18 |
| **MCF-7** |  |  |  |  |  |
| Unprocessed | 2.5x10<sup>6</sup> | 3420 | 9.5 | 100 | 25.77 |
| Stained | 2.5x10<sup>6</sup> | 3776 | 9.6 | 100 | 14.18 |
| Unprocessed | 1.5x10<sup>6</sup> | 6240 | - | 50 | 10.05 |
| Stained | 1.5x10<sup>6</sup> | 2712 | - | 50 | 3.14 |
| Cyclin B1 neg | 100000 | 279 | 9.1 | 100 | 14.99 |
| Cyclin B1 pos | 93000 | 334.5 | 8.8 | 100 | 30.11 |
| Cyclin B1 neg | 200000 | 1290 | 8.9 | 100 | 35.61 |
| Cyclin B1 pos | 131000 | 366 | 9.1 | 100 | 38.65 |

We then generated RNA-seq libraries from each of these RNA samples using a NEBNext Ultra II Directional RNA Library Prep kit with Poly(A)+ RNA Purification Module according to manufacturers’ instructions. RNA inputs and final library yields are given in Table 1. Equivalent PCR cycle numbers were used for library amplification between pairs of unprocessed and stained samples, and library yields obtained were no more variable than we would normally expect between samples. This shows that RNA recovered from glyoxal-fixed cells does not retain sufficient glyoxal adducts to impair RNA binding to oligo(dT) or RNA library construction. Log2-transformed read-counts for all annotated genes in the directly harvested compared to the fixed and stained samples correlated well (Fig. 3B, R^2^=0.989 for MCF-7 and R^2^=0.991 for COLO205), showing no major systematic change in mRNA distribution. Gene expression analysis between the sets of directly harvested versus fixed and stained cells by DEseq2 (p<0.05) identified 2 and 61 differentially expressed genes in MCF7 and COLO205 datasets respectively out of 60099 genes annotated in Ensembl GRCh38, indicating that very few genes are systematically affected by the staining and purification procedure, and the 61 genes did not significantly enrich for any functional category by GO analysis.

Finally, we performed two replicate experiments in which MCF-7 cells were stained and flow sorted into CCNB1 positive and negative fractions. Populations were gated for single cells, then gates for CCNB1 positive and negative populations were defined based on a control sample that was incubated with the antibody diluent, without adding the primary antibody (Fig. 3C). Frequency distributions of DAPI intensity in the unsorted cells versus the CCNB1 +ve and-ve populations showed that CCNB1 +ve cells were highly enriched for 2n DNA content as expected (Fig. 3D). 1-2×10^5^ cells of each fraction were harvested, RNA was extracted using TRI Reagent, analysed on a Bioanalyzer, and mRNA-seq libraries prepared and sequenced as above. The correlation between the two biological replicates of CCNB1-positive cells was excellent (Fig. 3E top, R^2^=0.992), whilst the CCNB1-negative fractions were slightly more divergent indicating differences between the cell cultures, although the correlation was still very strong (Fig. 3E bottom, R^2^=0.988). DEseq2 analysis (p<0.05) identified 161 significantly different expressed genes between CCNB1-positive and CCNB1-negative samples (Fig. 3F) including *CCNB1* along with many other known cell-cycle regulated transcripts (Table S1). Unsurprisingly, GO enrichment analysis for these transcripts identifies “mitotic cell cycle process” as the most enriched category (FDR corrected q-value 2.7×10^-65^) along with ~300 other significant enrichments (q<0.05) almost all of which are directly connected to cell cycle progression (Table S2). An equivalent experiment performed with COLO205 cells will be described elsewhere.

Overall, these data show that cells can be stained for an intracellular antigen and accurately sorted using our glyoxal fixation protocol, yielding high-quality, largely unbiased RNA samples that give rise to high quality RNA-seq libraries.

## Discussion

Although formaldehyde and glyoxal can be utilised similarly in both protein fixation and RNA denaturation, they react very differently with mixed RNA-protein substrates. This chemical difference gave rise to our prediction that glyoxal should fix cell samples without permanently modifying or crosslinking RNA. Here we have confirmed that RNA remains intact and accessible after glyoxal fixation, even through extended staining and sorting procedures. We further show that mRNA recovered from glyoxal-fixed cells can be purified by hybridisation to oligo(dT) and can be efficiently reverse transcribed.

Glyoxal fixative is easy to prepare and use, and should be straightforward to substitute for formaldehyde in most protocols. However we find that staining protocols do need to be modified to maintain RNA quality: firstly, inclusion of RNasin or an equivalent placental RNase inhibitor proved to be important. Secondly, we performed antibody incubations on ice to further reduce RNase activity, though we suspect this is not absolutely required. Thirdly, some staining reagents are clearly incompatible with RNA isolation - harsh detergents such as Triton X-100 allow RNA to escape the cell (an unavoidable consequence of the RNA not being crosslinked), and serum can contain RNase although RNA compatible blocking agents are commercially available [8]. We found it beneficial to perform trial stainings on the sample-type of interest and assess RNA quality by Bioanalyzer or RNA mini-gel (for which we provide a simple and robust protocol in Materials and Methods) as well as confirming staining quality before attempting a cell sorting experiment. It should however be noted that we did not need to render bulk staining solutions RNase free (e.g.: using DEPC), nor did we undertake any special cleaning of the flow cytometer or use special sheath fluid that would make routine use of these methods more arduous.

Whether glyoxal is a better or worse fixative than formaldehyde for microscopy studies of protein antigens remains a matter of dispute, and likely varies depending on precise cell type and target [27, 33]. In contrast, our data confirm that glyoxal is a far superior fixative for RNA applications because RNA remains extractable and not permanently modified in fixed cell samples. This means that substitution of formaldehyde with glyoxal and minor adjustments to staining buffers should be sufficient to render standard cell staining and sorting protocols compatible with a wide range of RNA methods including bulk and single cell RNA-seq.

## Materials and Methods

Step-by-step up-to-date protocols describing these methods are available from the JH group website at https://www.babraham.ac.uk/our-research/epigenetics/jon-houseley/protocols

### Cells and cell culture

COLO205 and MCF7 cell lines were provided by the laboratory of Dr. Simon J Cook at the Babraham Institute. All cell culture reagents were purchased from ThermoFisher Scientific, UK. Cells were cultured in RPMI-1640 (#21875) (COLO205) or Dulbecco’s modified Eagle’s medium (DMEM) (#41966) (MCF-7) supplemented with 10% (v/v) foetal bovine serum (#10270), penicillin (100 U mL-1), streptomycin (100 mg mL-1) (#15140) and 2 mM glutamine (#25030) in a humidified incubator at 37°C and 5% (v/v) CO_2_.

Cell line identity was validated based on RNA-seq data generated in this work using Cell Line Sleuth, developed by Simon Andrews of the Babraham Institute Bioinformatics Facility (https://github.com/s-andrews/celllinesleuth)

### RNA extraction

Sorted cells were pelleted by centrifugation at 2000 x *g* for 3 minutes at 4°C. For RNA isolation from cells grown in monolayers, cells were trypsinized, centrifuged at 300 x *g* for 5 minutes and washed once with PBS. Cell pellets were lysed in 1 mL TRI reagent (Sigma, T9424) and mixed until a homogeneous lysate was obtained. For phase separation, 0.2 mL chloroform was added to the lysate, mixed thoroughly and allowed to stand for 10 minutes prior to centrifugation at 12,000 × *g* for 15 min at 4°C. The upper aqueous phase containing RNA was transferred to a fresh tube, and 0.5 mL 2-propanol per mL original lysate was then added, samples mixed and allowed to stand for 10 minutes. Samples were centrifuged (12,000 × *g* for 10 min at 4°C), and RNA pellets washed with 1 mL 75% (v/v) ethanol (7,500 × *g* for 5 min at 4°C). Pellets were air-dried for 5-10 minutes and resuspended in RNase-free water.

RNA extraction from cells using Qiagen’s RNeasy extraction kit (Qiagen, 74104) was carried out according to the manufacturer’s instructions. Briefly, 1.5 x 10^6^ cells were lysed in a 350 μl cell lysis buffer and the lysate was mixed with equal amounts of 70% ethanol. The resulting mixture was applied to the RNeasy silica membrane and centrifuged (> 8000 x *g* for 15 seconds) to allow binding of RNA to the membrane. The membrane was washed twice by centrifugation (> 8000 x *g* for 15 seconds) with wash buffers to eliminate contaminants. RNA was eluted in 30μl nuclease-free water.

### RT-qPCR

Total RNA (100 ng) from COLO205 cells was subjected to one-step RT-qPCR analysis using the Luna Universal One-Step RT-qPCR Kit (NEB, E3005S) according to the manufacturer’s instructions. For each reaction, amplification was carried out in 20 μl reaction mixture containing 5 μl RNA sample (20 ng/μl), 10 μl Luna Universal One-Step Reaction Mix (1x), 1 μl Luna WarmStart RT Enzyme Mix (1x), 0.8 μl each of forward and reverse primers (0.4 μM) and 2.4 μl nuclease-free water. The PCR cycling conditions were reverse transcription at 55°C for 10 minutes, initial denaturation at 95°C for 1 minute, followed by 40 cycles of denaturation at 95°C for 10 seconds and extension at 60°C for 30 seconds on the Bio-Rad CFX-96 qPCR machine (Bio-Rad, UK). Data were analysed with CFX-manager software v.3.1. Details of primer sequences are provided in Table S3, amplicons were longer than normal in qPCR to better reflect the size range of cDNA fragments used in RNA-seq library construction.

### RNase-free technique

Filter tips and RNase-free tubes were used throughout this project. Certified RNase-free water (Sigma W3513) was used for elution and storage of RNA. Bulk solutions (PBS for staining, BPTE for electrophoresis, etc.) were assembled from standard laboratory chemicals and deionised or milliQ water without further purification. DEPC treatment was not used at any point. Gel electrophoresis was performed in a dedicated mini-gel tank although we consider this optional.

### RNA mini-gels

This is a very simple and reliable RNA gel electrophoresis protocol, a minor adjustment of the method originally described here [34]:

Assemble 10x BPTE buffer: 30 g PIPES free acid (Melford P40140), 60 g Bis-Tris (Melford B75000), 20 ml 0.5 M EDTA pH8, deionised water to 1L, stir or shake to dissolve. Store at room temperature >1 year, dilute to 1x with deionised water before use.

Assemble RNA denaturation mix: 6 ml DMSO (Sigma 34869), 2 ml 40% glyoxal solution (Sigma 50649, we do not deionise this), 1.2 ml 10x BPTE, 0.6 ml 80% glycerol. Split in 1 ml aliquots, keep a working aliquot at −20°C that can be used many times, store remaining aliquots at −70°C.

Obtain ethidium bromide solution (10 mg/ml, Bio-Rad 1610433), ensure this is less than 2 years old. Ideally keep frozen aliquots for RNA gels. Problems with migration, particularly the molecular weight marker, are most frequently due to aged ethidium solutions in our experience.

Add ethidium bromide solution at a final concentration of 50 μg/ml to sufficient RNA denaturation mix for 15 μl per sample including DNA molecular weight marker, vortex well (solution should be pink).

Add up to 3 μl RNA to 15 μl RNA denaturation mix containing ethidium bromide and incubate 1-2 hours at 55 °C. Treat the molecular weight marker in the same manner.

Cast a 1.2% agarose gel in 1x BPTE, and assemble apparatus with 1x BPTE as the running buffer.

Remove samples from heating block and load onto gel within 15 minutes.

Run gel at 90 V on a standard mini-gel system for 30 minutes to 1 hour before imaging.

### Preparation of glyoxal fixation solution

Glyoxal was purchased as a 40% aqueous solution from Sigma (50649), stored at 4°C, checked for precipitation before use and if necessary warmed to 55°C until precipitate re-dissolved. This solution was not deionised at any point. A 3% glyoxal solution at pH 4-5 was used in all experiments as described [27]. The solution was prepared by mixing 2.8 ml water, 0.79 ml 100% ethanol, 0.31 ml 40% glyoxal and 30 μl acetic acid. The pH of the solution was adjusted to 4-5 value (determined by pH paper) with a few drops of 1 M NaOH.

### Glyoxal fixation and staining

Trypsinise cells and collect the cell pellet in a 1.5 ml tube.

Wash cells once with 1 ml PBS and remove the supernatant.

Gently re-suspend cells in 100 μl of 3% Glyoxal fixation solution with 1:25 RNasin Plus (Promega N261B) and incubate for 15 minutes on ice.

Add 1 ml of 1xPBS with 1:100 RNasin Plus and centrifuge at 2000 x *g* for 3 minutes at 4°C. Discard the supernatant.

Gently re-suspend cells in 100 μl of ice-cold methanol (100%) with 1:25 RNasin Plus slowly (drop by drop while gently vortexing cells) to cells and incubate on ice for 30 minutes.

Add 1 ml of 1% BSA in PBS with 1:100 RNasin Plus and centrifuge at 2000 x *g* for 3 minutes at 4°C. Discard the supernatant.

Gently re-suspend cells in 100 μl of diluted antibody (in this case anti-CCNB1, CST 12231S, diluted 1:200) in 1% BSA in PBS with 1:25 RNasin Plus and incubate for 1 hour on ice.

Add 1ml of 1% BSA in PBS with 1:100 RNasin Plus and centrifuge at 2000 x *g* for 3 minutes at 4°C. Discard the supernatant.

Re-suspend cells in 100 μl of diluted secondary antibody (in this case Alexa Fluor 488 donkey anti-rabbit antibody, ThermoFisher Scientific A21206, diluted 1:1000) in 1% BSA in PBS with 1:25 RNasin Plus. Incubate on ice for 30 minutes in the dark.

Add 1 ml of 1% BSA in PBS with 1:100 RNasin Plus and centrifuge at 2000 x *g* for 3 minutes at 4°C. Discard the supernatant.

Re-suspend the cell pellet in 200 μl of 1% BSA in PBS with 1:100 RNasin Plus.

### Flow cytometry

Prior to sorting, all samples were filtered to eliminate cell aggregates by passage through a sterile, 30 μm CELLTrics filter (Sysmex, 04-004-2326) into 5 mL polypropylene round-bottom tubes (Scientific laboratory supplies, 352063). Samples were then labelled with 4,6-diamidino-2-phenylindole (DAPI) (Sigma, D9542) at a final concentration of 1 μg/mL in PBS. Cells were sorted using a 100 μm nozzle (at 20PSI) on a BD FACSARIA III cell sorter (BD Biosciences, UK) with cooling of the sample and collection chambers enabled. Cells were collected in 1.5 mL RNase-free tubes pre-coated with 1% BSA in PBS supplemented with 1:25 RNasin Plus RNase inhibitor. The Alexa488 was excited by a 488nm laser and 530/30 bandpass filter used for collection of fluorescence. The DAPI was excited by the 355nm laser and its emission collected using a 450/50 filter.

### Preparation and sequencing of RNA-seq libraries

RNA prepared from flow sorted cells was quality controlled on a Bioanalyzer 2100 (Agilent) using RNA Pico 6000 chips (Agilent 5067-1513). RNA-seq libraries were prepared using the NEBNext Ultra II Directional RNA Library Prep Kit for Illumina (NEB E7760S) with the NEBNext Poly(A) mRNA Magnetic Isolation Module (NEB E7490S), following the protocol provided by the manufacturer with the exception that two successive 0.9x AMPure purifications were performed on the final amplified libraries. All libraries were amplified using 12 PCR cycles. Library quality was assessed on a Bioanalyzer 2100 using High Sensitivity DNA chips (Agilent 5067-4626) and quantification performed with a KAPA Library Quantification Kit (Roche, KK4844). Libraries were sequenced by the Babraham Institute Next Generation Sequencing Facility on an Illumina HiSeq2500 in Rapid Run 50bp Single End mode.

### Analysis of RNA-seq data

After adapter and quality trimming using Trim Galore (v0.6.2), RNA-seq data were mapped to human genome GHCh38 using HISAT2 v2.1.0 [35] by the Babraham Institute Bioinformatics Facility. Mapped data were imported into SeqMonk v1.47.0 (https://www.bioinformatics.babraham.ac.uk/projects/seqmonk/) and antisense reads mapping to each annotated gene were quantified and normalised to the total read count in each library. Scatterplots and hierarchical clustering were generated using SeqMonk, and analysis of differential gene expression performed using DEseq2 via SeqMonk [36]. GO analysis was performed using GOrilla (http://cbl-gorilla.cs.technion.ac.il/) [37, 38]. Quoted q-values for GO analysis are FDR-corrected according to the Benjamini and Hochberg method (q-values from the GOrilla output) [39].

All RNA-seq data has been deposited at GEO under accession number GSE158177.

## Acknowledgements

We would like to thank Rachael Walker and Rebecca Roberts of the Babraham Institute Flow Cytometry facility for help with setting flow sorting, the Babraham Institute Next Generation Sequencing and Bioinformatics facilities for generating and processing RNA-seq data. We particularly appreciate the effort invested by Simon Andrews of the Babraham Institute Bioinformatics facility for developing Cell Line Sleuth. We also thank Jonathan Clark for advice on chemical activities of cross linkers, Simon Cook and lab for cell lines, and Rachael Walker and Stephen Clark for critical reading of the manuscript.

